# The Interaction Between Miro and TRAK is not Required for Bulk Mitochondrial Trafficking

**DOI:** 10.64898/2026.05.01.722185

**Authors:** Christian Covill-Cooke, Milli Owens, Andreas Prokop, Benoît Kornmann

## Abstract

In metazoans, mitochondria optimally distribute to sites of need through long-range transport events on microtubules. The prevailing model for this trafficking mechanism is that the tail-anchored calcium-binding GTPase, Miro, recruits cytosolic TRAK and associated molecular motors to the outer mitochondrial membrane. Therefore, Miro is proposed to be an obligate adaptor for TRAK required for bulk mitochondrial transport, a process that is considered particularly important for long-range trafficking in neurons, and thus, for viability. Here, we impaired Miro-TRAK interaction *in vivo* by introducing a point mutation into the *Drosophila* TRAK orthologue Milton, that impairs its interaction with Miro, based on recent structural evidence. Flies harbouring this point mutation are viable to adulthood. Moreover, neurons carrying this mutation exhibit little to no observable reduction in axonal mitochondria. Mutant flies, however, display progressive loss of motor function with age and reduced lifespan. We therefore call into question the long-standing view that Miro plays an obligatory role in mitochondrial trafficking and challenge the canonical model for mitochondrial transport.

## Introduction

Finite pools of subcellular components must be distributed throughout the cell to support cellular functions. In metazoans, this is achieved in large part by long-range trafficking on microtubules by the molecular motors, kinesins and dynein (Barlan and Gelfand, 2017). One of the most thoroughly described mechanisms of microtubule-dependent organelle transport is mitochondrial trafficking, which predominantly occurs via the recruitment and activation of kinesin-1 or dynein at the outer mitochondrial membrane (OMM) through interaction with Trafficking Kinesin Proteins (TRAK1/2 in humans and Milton in *Drosophila*, collectively referred to as TRAKs from hereon) (Glater et al., 2006; van Spronsen et al., 2013; Stowers et al., 2002; Canty et al., 2023; Henrichs et al., 2020; Fenton et al., 2021). As TRAKs are soluble proteins, they must bind to proteins at the OMM. In the current prevailing and widely accepted model, the Mitochondrial Rho GTPases (MIRO1 and 2 in mammals, Miro in *Drosophila*) function as the TRAK adaptors at the OMM (Glater et al., 2006; Fransson et al., 2006); these three components (molecular motors, TRAK and Miro) are therefore considered the core trafficking machinery. The biochemical features of Miro, namely a calcium binding GTPase, has positioned Miro as an obvious avenue not only for promoting transport, but also regulating mitochondrial position in response to cellular cues, e.g., halting mitochondria movement upon increased cytosolic calcium levels, presumably by disengaging the trafficking machinery (Macaskill et al., 2009; Wang and Schwarz, 2009; Saotome et al., 2008). Support for the canonical model comes from two main observations: i) The members of this core trafficking machinery directly interact, i.e. TRAK directly binds to Miro and kinesin (Glater et al., 2006; Macaskill et al., 2009; MacAskill et al., 2009; Wang and Schwarz, 2009; Stowers et al., 2002; Ravitch et al., 2025; Covill-Cooke et al., 2024; van Spronsen et al., 2013); ii) Loss of TRAK or Miro largely phenocopy each other. In *Drosophila*, this phenocopy manifests as a dramatic loss of mitochondria from motoneuronal axons and larval lethality of *milton* (*milt*) and *Miro* null mutants (Stowers et al., 2002; Guo et al., 2005; Russo et al., 2009). In mammalian cultured cells and neurons, loss of Miro leads to profound changes in mitochondrial distribution (Kanfer et al., 2015; López-Doménech et al., 2018, 2016), and specific loss of *Miro1* in mice cause perinatal death through insufficient innervation of the lungs, and depletion of mitochondria in the axons of the cerebrospinal tract (López-Doménech et al., 2018, 2021; Nguyen et al., 2014).

The model above has been ubiquitous to the mitochondrial trafficking field since its formulation in the 2000s, and underpins most – if not all – of the work on the regulation of mitochondrial trafficking in different cellular contexts (Macaskill et al., 2009; Wang and Schwarz, 2009; Wang et al., 2011; Ashrafi et al., 2014; McKenna et al., 2025). It also serves as a broader paradigm for organelle transport (Barlan and Gelfand, 2017).

However, scattered observations are at odds with this framework, specifically on the role of Miro in mitochondrial transport. Firstly, although Miro is the only complex component with a transmembrane domain, TRAK1/2 and molecular motors can localise to mitochondria in its absence (López-Doménech et al., 2018; Mitchell et al., 2025). This Miro-independent mitochondrial recruitment is achieved, at least in part, by features within the C-terminus of TRAKs (MacAskill et al., 2009; Glater et al., 2006; Mitchell et al., 2025). Secondly, whilst loss of Miro causes large changes to mitochondrial distribution, it does not completely block microtubule-dependent transport in cultured mammalian cells (López-Doménech et al., 2018) and results in a less severe mitochondrial phenotype than loss of *milt* in cultured primary *Drosophila* neurons (Liew et al., 2026). Therefore, the broad phenocopy of *milt* and *Miro* null *Drosophila* mutants, is not as complete as would be expected from the model. Finally, various direct interactors of Miro have been identified in processes as diverse as lipid transport and mitophagy (Covill-Cooke et al., 2024; Guillén-Samander et al., 2021; Oeding et al., 2018; López-Doménech et al., 2018; Koszela et al., 2025; Kanfer et al., 2015; Kornmann et al., 2011), raising the possibility that Miro knockout perturbs multiple aspects of mitochondrial biology. Consequently, changes to mitochondrial distribution may arise indirectly from disruptions to these processes, confounding interpretation of Miro’s role in mitochondrial trafficking. Thus, only by selectively disrupting Miro-TRAK interaction, leaving other functions of both proteins intact, can the role of Miro in TRAK-dependent mitochondrial transport be rigorously defined.

We and others have recently described a conserved interface for the Miro-TRAK interaction, whereby a disordered loop in TRAK interacts with a conserved surface on Miro formed by its first GTPase and calcium-binding EF hand with LM helices (ELM1) domains. An essential feature of this interaction is the insertion of a leucine residue of TRAK’s Miro-binding domain into a deep hydrophobic cavity of the ELM1 domain in Miro (Figure 1A). Mutation of this leucine impairs Miro-TRAK interaction (Ravitch et al., 2025; Covill-Cooke et al., 2024).

**Figure 1:**
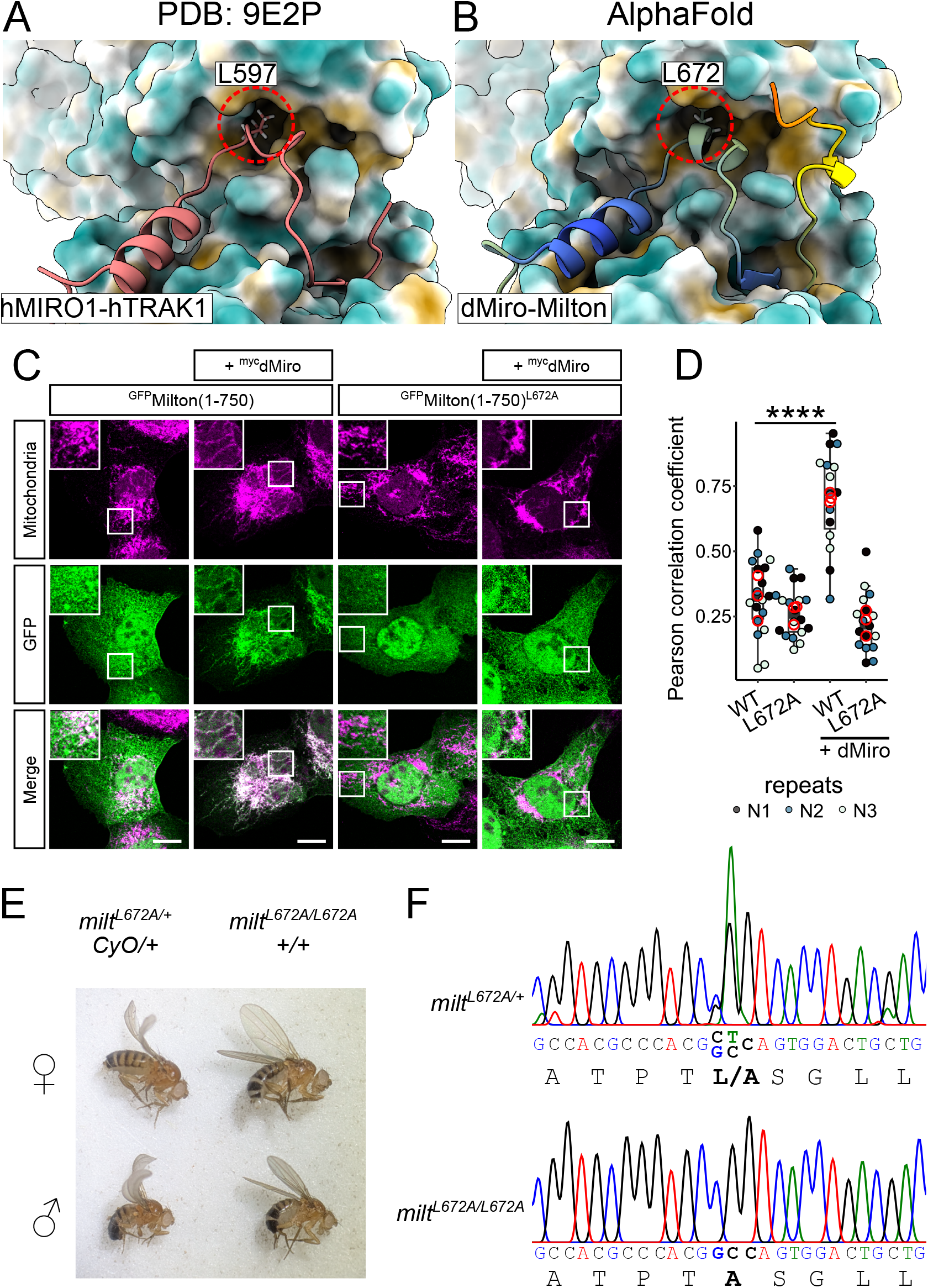
Milton^L672A^ prevents the interaction with Miro but does not lead to larval lethality. A) CryoEM structure from Ravitch *et al*. (2025) of the human TRAK1 (shown as red ribbon) interacting with MIRO1 (shown as surface model). Red circle highlights Leucine 597 essential for the interaction. B) AlphaFold3 prediction of the corresponding *Drosophila* protein domains, where the cognate leucine is in position 672. C) Representative images of U2OS cells expressing the *Drosophila* proteins Miro together with GFP-tagged variants Milton(1-750) or Milton(1-750)^L672A^. Mitochondria are stained for TOMM20 (magenta). Scale bar is 10 µm. D) Quantification of the experiments shown in C determining the mitochondria enrichment of Milton(1-750) and Milton(1-750)^L672A^ by Pearson’s coefficient (correlation of GFP signal with mitochondrial marker TOMM20). Statistical testing was performed using two-way ANOVA with Tukey’s posthoc test on the average coefficients of the three independent repeats, with **** denoting p<0.0001. E) Representative images highlighting adult viability upon the loss of CyO and therefore of the homozygous *milt*^*L672A*^ flies. F) Sanger sequencing chromatograms to confirm the genotype of the *milt/milt*^*L672A*^ and *milt*^*L672A*^*/milt*^*L672A*^ flies.

Given that the mitochondrial trafficking field, including the discovery of the Miro-TRAK interaction, is built upon seminal work in fruit flies, where core components of the mitochondrial trafficking machinery are conserved as single orthologues (Guo et al., 2005; Stowers et al., 2002; Glater et al., 2006), we used this system to test the functional importance of Miro-TRAK interface *in vivo*. We have generated a point mutant in the conserved leucine of *Drosophila* TRAK-orthologue Milton to impair the interaction with Miro, while preserving other functions of both proteins. As such, we can now test whether the Miro-TRAK interaction is truly required for bulk mitochondrial transport. Surprisingly, we find that the Miro-Milton interaction is dispensable for the bulk distribution of mitochondria in axons and for adult viability.

## Results

### Milton^L672^ is required for binding to Miro

Single point mutants have previously been identified in the mammalian system which impair binding between MIRO and TRAK *in vitro*, in cultured cells and in the yeast-two-hybrid assay (Ravitch et al., 2025; Covill-Cooke et al., 2024). As the Miro binding motif of TRAK is highly conserved (Covill-Cooke et al., 2024), including the leucine residue that inserts into Miro’s hydrophobic pocket, we mutated the corresponding leucine 672 in the *D. melanogaster* protein (Milton) to alanine, to impair its interaction with Miro (Figure 1A-B). We note that a second Miro-binding site within the human TRAK1/2 proteins has been proposed (Baltrusaitis et al., 2023; Ravitch et al., 2025); however, this site is not conserved outside of vertebrates and, crucially, is missing in *Drosophila*. To confirm that the L672A substitution in Milton prevents binding to *Drosophila* Miro, as is the case for human TRAK/MIRO, we employed a strategy successfully applied previously (Glater et al., 2006); a cytosolic C-terminally truncated form of Milton, Milton(1-750), lacking Miro-independent mitochondrial targeting features, is only recruited to mitochondria upon *Drosophila* Miro overexpression, which therefore provides a readout of the Miro-Milton interaction (Glater et al., 2006). We found that while wild-type Milton(1-750) was robustly recruited to mitochondria upon *Drosophila* Miro co-expression, Milton(1-750)^L672A^ remained cytosolic in the same conditions (Figure 1C-D). Absence of recruitment of Milton(1-750)^L672A^ was also observed in cells with high Miro expression levels which are known to induce perinuclear collapse of the mitochondrial network (Supp. Figure 1) (Glater et al., 2006). Therefore, the Milton^L672A^ mutant impairs the interaction between Miro and Milton, as predicted.

### Milt^L672A^ homozygous flies are viable

Loss of *milt* or *Miro* results in larval death (Guo et al., 2005; Stowers et al., 2002). This phenocopy of the null alleles is an important part of the data supporting a model where Miro and Milton make up the essential core trafficking complex. To test whether the lethality of *Miro* mutants results specifically from loss of Milton-dependent trafficking rather than other Milton-independent processes, we used CRISPR-Cas9 editing to engineer the L672A mutation into the endogenous *milt* locus of flies. Four independent heterozygous *milt*^*L672A*^ mutant fly lines were obtained and balanced with the CyO balancer chromosome. Crossing these CyO/*milt*^*L672A*^ lines yielded progeny lacking the balancer, indicating the presence of viable homozygous *milt*^*L672A/L672A*^ mutant flies, which was confirmed by Sanger sequencing (Figure 1E-F). These homozygous lines were stably maintained for many generations. Since impairing Miro-Milton interaction does not recapitulate the phenotypes of *Miro* null mutants, we conclude that, the larval lethality observed in *Miro* flies is not solely caused by defective Milton-dependent mitochondrial transport.

### The Miro-Milton interaction is not essential for mitochondrial distribution

Both Miro and Milton are necessary for mitochondria distribution in axons. We thus sought to assess the effect of disrupting Miro-Milton interaction on neuronal mitochondrial distribution. To achieve this, we generated primary neuron cultures from wild-type and homozygous *milt*^*L672A*^ mutant embryos and quantified the ability of mitochondria to distribute from the somata to axonal processes. As controls, we used previously characterised *milt*^*92*^ (Stowers et al., 2002) and *Miro*^*sd32*^ null mutant neurons (Guo et al., 2005). These recapitulated the reported accumulation of mitochondria in the cell bodies, and their near absence in axons (Figure 2A-C), with a milder phenotype in *Miro*^*sd32*^ (Liew et al., 2026). The distribution of mitochondria in homozygous *milt*^*L672A*^ neurons and hemizygotic heteroallelic *milt*^*L672A*^*/milt*^*92*^ neurons, however, was comparable to wild-type neurons, with mitochondria present throughout the axonal processes (Figure 2A-C; Supp. Figure 2A-C). Therefore, bulk microtubule-dependent trafficking of mitochondria does not require Miro-Milton interaction.

**Figure 2:**
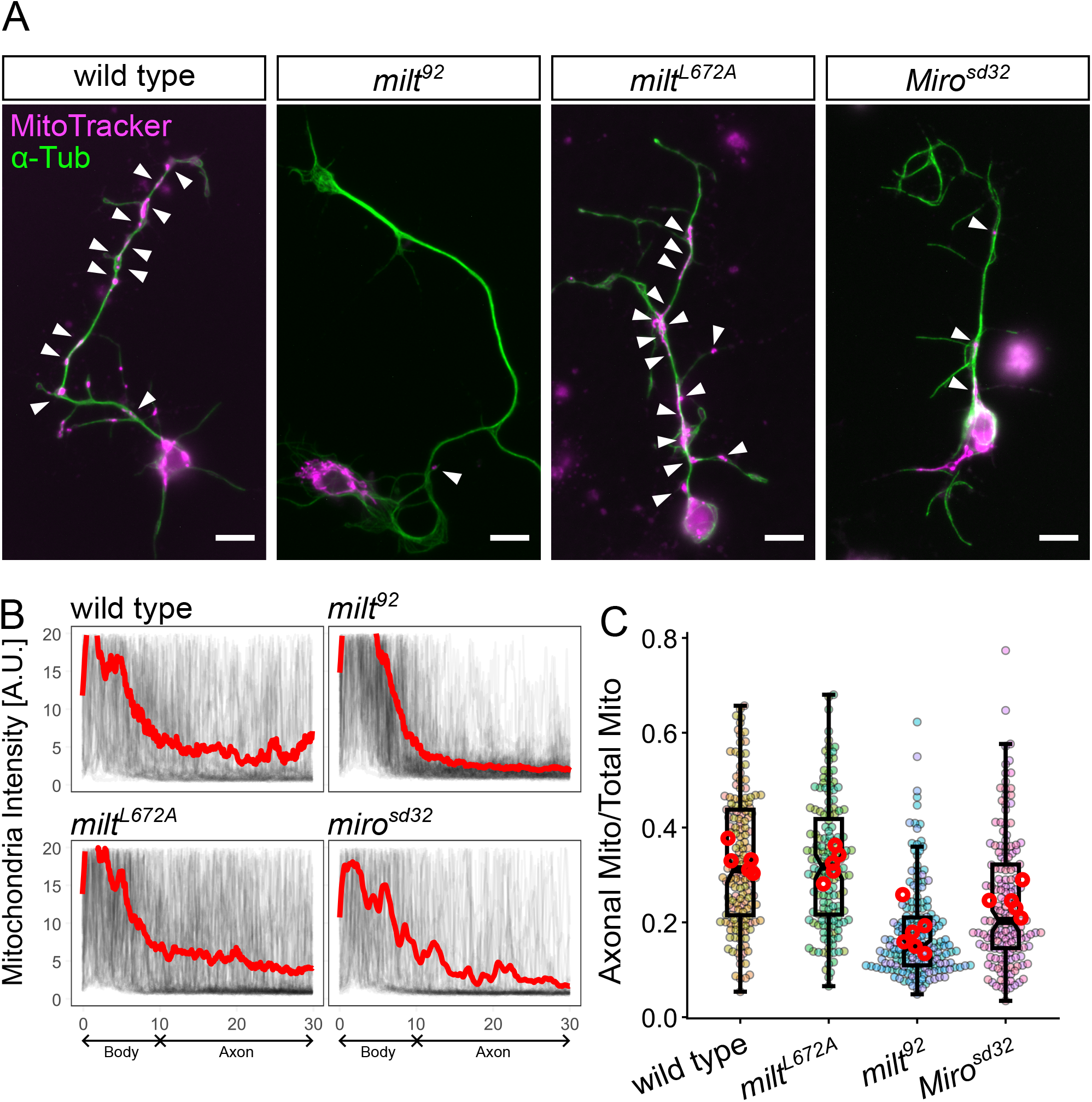
Abolishing the Miro-Milton does not impact mitochondrial distribution. A) Representative images of wild type, *milt* null *Miro* null and homozygous *milt*^*L672A*^ neurons. Green shows microtubules (anti-α-tubulin) and magenta mitochondria (MitoTracker). Scale bar is 10 µm. B) Overlay of the mitochondrial intensity distributions in axons of wild type, *milt* null *Miro* null and homozygous *milt*^*L672A*^ neurons. Grey lines represent individual cells and the red line indicates the average distribution for each genotype. C) Quantification of the axonal mitochondrial content defined as the fraction of total mitochondrial intensity located between 12 µm and 30 µm from the soma. Number of cells analysed in B and C, WT, 151 cells; *milt*^*L672A*^, 152 cells; *milt*^*92*^, 180 cells; *Miro*^*sd32*^, 158 cells. Five replicates per genotype (indicated by colour shades). Statistical significance was assessed using the pairwise Wilcoxon rank-sum test: WT and *milt*^*L672A*^ differ from both *milt*^*92*^ and *Miro*^*sd32*^ *(p* < 0.01 and *p* < 0.05, respectively) whereas no significant difference is found between WT and *milt*^*L672A*^.

### Miro-Milton interaction is required in ageing

We observed that, while heterozygotes flies expressing one single copy of wild-type *milt* (*milt*^*+*^*/milt*^*92*^) are viable (Stowers et al., 2002), heteroallelic flies expressing one single copy of *milt* L672A (*milt*^*L672A*^*/milt*^*92*^) failed to develop to adulthood. This suggests that, although dispensable for bulk mitochondrial trafficking, the Miro-Milton interaction nonetheless plays an important physiological role. Consistent with this interpretation, *milt*^*L672A*^ flies displayed a progressive loss of motor function, as assessed by negative geotaxis. No noticeable differences in climbing were observed when comparing freshly eclosed *milt*^*L672A*^ homozygous to heterozygous counterparts (Figure 3A; Supp. Video 1), but by five weeks of age, *milt*^*L672A*^ homozygous flies showed an almost complete lack of climbing (Figure 3A; Supp. Video 2), coinciding with early mortality (Figure 3B).

**Figure 3:**
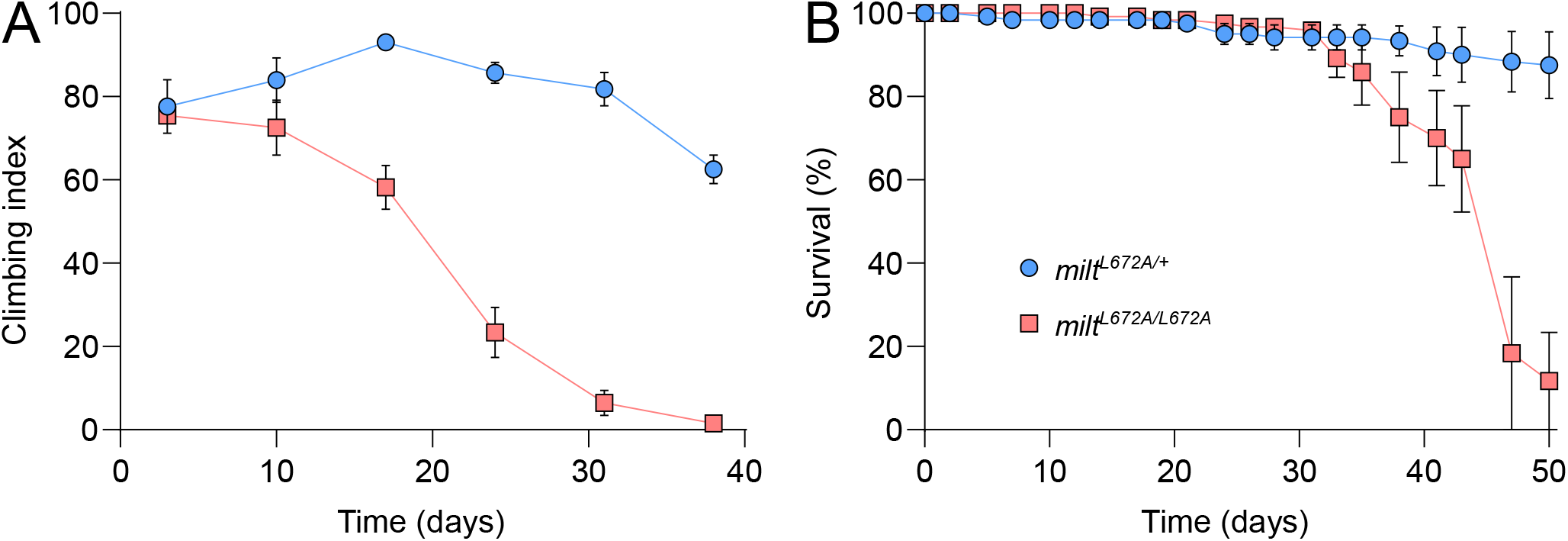
Milt^L672A^ homozygous flies have an age-dependent loss of motor function and reduced lifespan. A) Quantification of percentage of flies that climb 3 cm within 4 seconds (climbing index) in *milt*^*L672A/+*^ heterozygous and *milt*^*L672/L672A*^ homozygous flies across 5 weeks. N=3 independent fly lines used. B) Quantification of survival the flies used in A over 50 days.

Therefore, the L672 residue of Milton required for Miro interaction is necessary for vitality and viability in adults. To conclude, impairing Miro-Milton interaction results in viable adult flies but appears to lead to progressive loss of motor function and reduced lifespan.

## Discussion

The interaction between Miro and Milton has long been thought to be an obligatory core feature of the mitochondrial trafficking machinery. Here, we show that perturbing this interaction does not affect the bulk distribution of mitochondria by Milton- and kinesin-dependent transport on microtubules.

### Miro, TRAK and mitochondrial trafficking

The identification of TRAK as the first known direct interactor of Miro (Glater et al., 2006; Giot et al., 2003), and the profound mitochondrial distribution defects observed upon loss of Miro (Guo et al., 2005; Nguyen et al., 2014; López-Doménech et al., 2018, 2016; Kanfer et al., 2015), have made Miro synonymous with microtubule-dependent mitochondrial transport. Evidence inconsistent with this canonical model of mitochondrial trafficking has, however, slowly emerged. In mouse embryonic fibroblasts lacking Miro1 and Miro2, the transport machinery including TRAK, kinesin-1 and dynein localise to mitochondria as in wild-type cells (López-Doménech et al., 2018), owing to recently characterized features in TRAK C-terminus (Mitchell et al., 2025). In parallel, an ever-growing repertoire of Miro-binding proteins was unveiled, suggesting roles for Miro extending far beyond mitochondrial transport (Covill-Cooke et al., 2024; Oeding et al., 2018; López-Doménech et al., 2018; Guillén-Samander et al., 2021; McKenna et al., 2025). These observations called for a re-evaluation of the role of Miro-Milton/TRAK interaction in mitochondrial transport.

### The impact of Miro on mitochondrial distribution and homeostasis

We find that impairing the Miro-Milton interaction does not impact the broad distribution of mitochondria in neuronal axons, thus calling into questions one of the most thoroughly studied models for organelle transport. Although Milton/TRAK’s role remains firmly linked to mitochondrial transport, that of Miro requires re-evaluation. Albeit not essential for mitochondrial transport, the Miro-Milton/TRAK interaction is nonetheless conserved from *Drosophila* to man, and our data indicate physiological relevance for longer-term health and survival, as underscored by the progressive defect in motor activity and reduced lifespan exhibited by *milt*^*L672A*^ homozygous flies. A potential model is that, though not central to trafficking, Miro might be regulating mitochondrial position in response to cellular cues. One particularly intriguing observation is that, while heterozygosity for *milt* (*milt*^*92/+*^) shows no apparent haploinsufficiency with respect to development, the *milt*^*L672A*^/*milt*^*92*^ heteroallelic hemizygous flies fail to reach adulthood. A straightforward interpretation could be that Miro-bound (wild-type) Milton is somewhat more active than Miro-free (L672A) Milton. As a result, halving the gene dosage of the former may be tolerated, while that of the latter may not, although we cannot formally exclude that the L672A mutation affects Milton activity independently of Miro binding. The use of our mutant model may therefore help in deciphering the exact role of Miro in mitochondrial transport, and assess if and how Miro stimulates TRAK/Milton activity, and in which conditions.

Because of the historical connection between Miro and mitochondrial transport, newly-identified Miro interactors are often interpreted exclusively through the lens of mitochondrial trafficking. For example, Miro is a target of the ubiquitin E3-ligase Parkin during mitochondrial damage (Wang et al., 2011; Sarraf et al., 2013). The function of this interaction during mitochondrial damage was therefore thought to allow Miro degradation and halt mitochondrial movement for efficient capture by the autophagosome during mitophagy (Wang et al., 2011; Shlevkov et al., 2016; Ashrafi et al., 2014), without considering the effects that Parkin-mediated Miro degradation might have on other functions of mitochondria such as lipid supply (Guillén-Samander et al., 2021) or fission regulation (Covill-Cooke et al., 2024; McKenna et al., 2025). The growing number of functionally diverse Miro interactors (Covill-Cooke et al., 2024) has made this position increasingly difficult to hold and left Miro knockout studies increasingly difficult to interpret. Now with the characterisation of the Miro-TRAK interface (Ravitch et al., 2025; Covill-Cooke et al., 2024), it is possible to move away from complete loss of function studies to specifically study the role of Miro in mitochondrial transport.

An important question remains; why does deleting Miro prevent mitochondrial distribution to neurites and cause larval lethality if preventing its interaction with Milton does not? The most straightforward explanation is that Miro possesses activities outside of its involvement with Milton/TRAK. Miro appears to be present in the common ancestor of all eukaryotes; TRAK- and microtubule-mediated mitochondrial transport are metazoan-specific features, and therefore, more recently evolved. Many direct interactors have been identified for Miro in mammals; these include the lipid transporter VPS13D, the E3-ligase Parkin, the actin motor MYO19, the microtubule-binding protein CENP-F and the fission regulators MTFR1/2/1L (Covill-Cooke et al., 2024; Oeding et al., 2018; López-Doménech et al., 2018; Guillén-Samander et al., 2021; Kanfer et al., 2015; McKenna et al., 2025). Therefore, upon loss of Miro, the function of these other interaction partners is likely to be affected and could lead indirectly to the observed mitochondrial trafficking defects. Of note, the above alternative clients are typically not conserved in *Drosophila* (MTFRs, MYO19, CENP-F), and when they are (Parkin and VPS13D), their Miro interaction domain is not (Covill-Cooke et al., 2024). We therefore surmise that the dysfunction of (an) alternative client(s) might be responsible for the severe phenotypes of *Miro* mutants. Such alternative clients are yet-to-be-identified.

## Materials and Methods

### Plasmids and antibodies

*Drosophila* cDNAs were obtained from the *Drosophila* Genome Resource Centre - Milton: LD33316 (DGRC Stock 5269), Miro: RE22983 (DGRC Stock 1095267). Residues 1-750 of Milton were cloned into pEGFP-C1 for N-terminal GFP-tagging and mammalian expression. Miro was cloned into pRK5-myc for N-terminal myc-tagging and mammalian expression. Mutation of L672A in ^GFP^Milton(1-750) was achieved by Quikchange.

Primary antibodies: anti-TOMM20 (rabbit; Santa Cruz, sc-11415, 1:500) and anti-myc (rabbit; Abcam, ab9106, 1:1000). Secondary antibodies: anti-mouse IgG H&L-AlexaFluor-647 (donkey; Abcam, ab1501017, 1:500), anti-rabbit IgG H&L-AlexaFluor-568 (donkey; Abcam, ab175470, 1:500).

### Fly stocks

Flies were maintained on standard cornmeal yeast extra media at 25°C with a 12-hour light–12-hour dark cycle. The following mutant stocks were used: *milt*^*92*^ (FBal0147011: 2bp deletion at aa248 introducing a frame shift; Tom Schwarz), *Miro*^*Sd32*^ (FBal0240382: 29 bp deletion frame-shifting at Y89; Tom Schwarz; FlyBase:FBal0240382), as well as balancer chromosomes: *CyO, P{GAL4-twi*.*G}2*.*2, P{UAS-2xEGFP}AH2*.*2* (RRID:BDSD_6662) and *TM3, P{GAL4-twi*.*G}2*.*3, P{UAS-2xEGFP}AH2*.*3, Sb*^*1*^ *Ser*^*1*^ (RRID:BDSD_6662).

### Generation of the Milt-L672A lines

Four *milt*^*L672A*^ lines were generated by Rainbow Transgenic Flies Inc, as previously (Gratz et al., 2014), by injecting the following into *y*,*w*; +/+; *Nos-Cas9-attp2*:

1. gRNA 5’ – TCCAGCAGTCCACTGAGCGT – 3’
2. Repair template 5’-AAACCCATGGAGGGATCGCAGACATTGCACCATTGGTC GCGCTTGGCCACGCCCACGgcCAGTGGACTGCTGGATGAGCGTCCCGGCGT AACGATCCGTGGTGGACGTGGCCTGGA -3’.

Clones positive for the *milt*^*L672A*^ mutation were balanced with CyO.

### Neuronal cultures

Egg lays were set the day before culturing by tipping the flies into a new vial with fly food and dry yeast. After 14-15 hours at 21°C, embryos were dechorionated in 50% sodium hypochlorite solution, collected in a sieve, selected under a fluorescent dissecting microscope for correct genotype (selected against green balancers) and the right stage (≥ stage 11) according to (Campos-Ortega and Hartenstein, 1997). After selection, 10-20 embryos per genotype were transferred to a 1.5 ml centrifuge tube and treated as described previously (Prokop et al., 2012). In brief, embryos were washed with 70% ethanol which was replaced by 100 µl Schneider’s medium. 100 µl of dispersion medium (2 ml of HBSS, 1 mg/ml Collagenase, and 2 mg/ml Dispase) was added and embryos were physically dispersed with a micro-pestle and incubated at 37°C for 5 mins. To slow down the reaction, 200 µl of Schneider’s medium was added, the solution was pipetted up and down and then centrifuged at 650 G for 4 mins. The solution was taken off, replaced with 90 µl Schneider’s medium which was pipetted up-and-down to disperse cell clumps. 30 µl were pipetted on three concanavalin A-coated coverslips, respectively. Coverslips were sealed with Vaseline onto special culture chambers. To make sure the cells stick to the coverslips, culture chambers were kept coverslip-down for 2 hrs and then turned coverslip-up (hanging-drop culture) to develop for 3 days at 26°C. Cells were cultured for 5 days before fixation.

### Immunocytochemistry and fluorescence microscopy

For Figure 1: U2OS cells were fixed with 4% PFA blocked for 30 min with 10% foetal bovine serum, 5 mg/ml bovine serum albumin and 0.2% Triton X-100 in PBS. Samples were stained with primary and secondary antibodies for 1 h each and imaged on a Zeiss LSM980 upright confocal microscope with a 40x objective (NA=1.3).

For Figure 2: *Drosophila* primary neurons were fixed for 30 min with 4% PFA in PBS. Cells were permeabilised 0.5% Tergitol (Sigma-Aldrich, Cat# 15S9) in PBS for 30 min and stained with primary and secondary antibodies for 2 h and 1.5 h, respectively. To visualise mitochondria, 30 µL of 400 nM MitoTracker Red CMXRos (Invitrogen; Cat# M7512) in Schneider’s Medium was applied to cells via culture media exchange and incubated for 30 min at 26°c prior to fixation. Imaging was performed using an AxioCam monochrome digital camera mounted on an Olympus BX50WI or BX51 compound fluorescent microscope.

### Image analysis

Figure 1: Colocalisation analysis of Milton(1-750) and mitochondria was performed on manually traced ROIs representing most of the cell to exclude nuclear signal. Pearson’s correlation coefficient was then calculated between the GFP and mitochondrial channels in R.

Figure 2: The longest axonal process was traced manually in ImageJ. Intensity profiles were then taken for the mitochondrial channel and plotted with distance from the centre of the soma. The ratio of mitochondrial signal between 7.5 µm and 30 µmover total signal was then obtained for each of the longest axons.

### Protein-protein interaction predictions

The interaction of full-length Milton (accession number: Q960V3-1) and Miro (Q8IMX7-1) were predicted using AlphaFold3 (Abramson et al., 2024), and visualised using ChimeraX (Pettersen et al., 2021).

### Climbing and survival assays

Forty *milt*^*L672A/+*^ heterozygous or forty *milt*^*L672A/L672A*^ homozygous flies of three independent lines were maintained with food change every 3 days. The number of dead flies was recorded every three days. For the climbing assay, videos were taken once per week of negative gravitaxis of flies after tapping the flies to the bottom of a tube. Climbing index was defined as the percentage of flies that climbed at least 3 cm in 4 seconds.

## Acknowledgments

The authors would like to thank Maximilian Fitz-James, and Petros Ligoxygakis and his lab for technical support and advice. We also gratefully acknowledge the Micron Advanced Bioimaging Facility (supported by Wellcome Strategic Awards 091911/B/10/Z and 107457/Z/15/Z) for their support & assistance in this work. This work was funded by the Wellcome Trust grant 214291/Z/18/Z (awarded to BK). The Manchester Fly Facility has been supported by funds from The University of Manchester and the Wellcome Trust (087742/Z/08/Z). We thank colleagues and the Bloomington Drosophila Stock Center (supported by NIH P40OD018537) for providing fly stocks as detailed in Materials and Methods.

## Supplementary Figure Legends

**Supplementary Figure 1:**
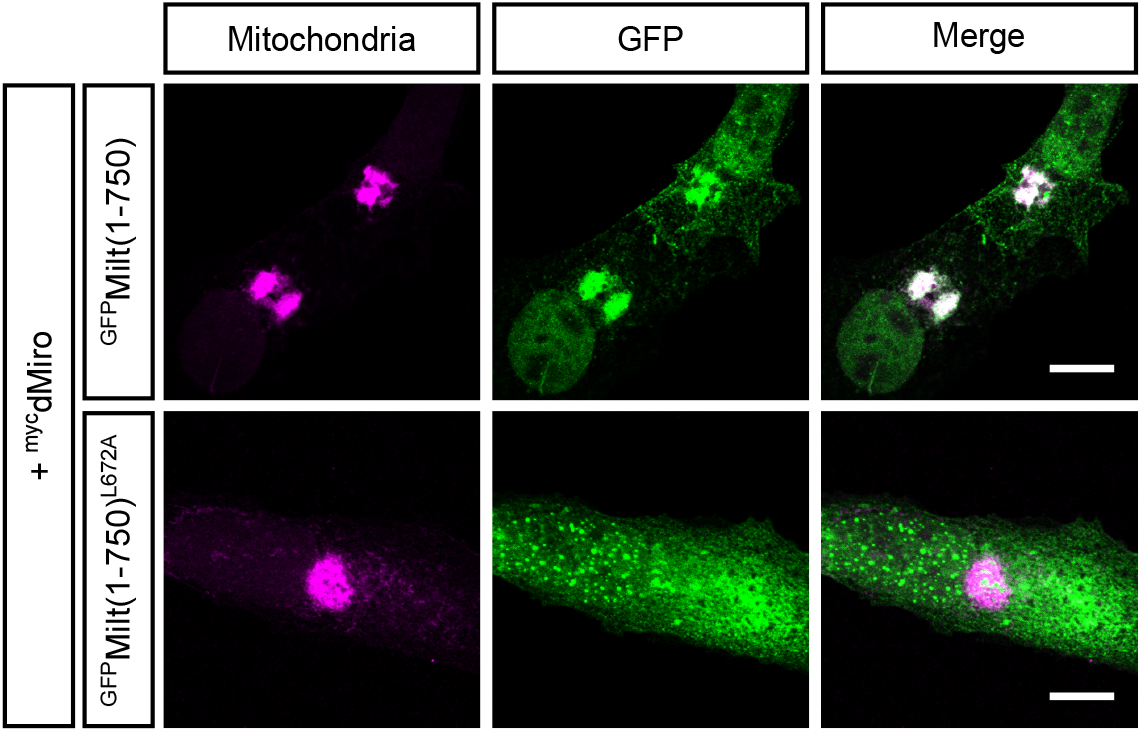
L672A mutation of Milton(1-750) does not localise to mitochondrial clustering induced by Miro overexpression.

**Supplementary Figure 2:**
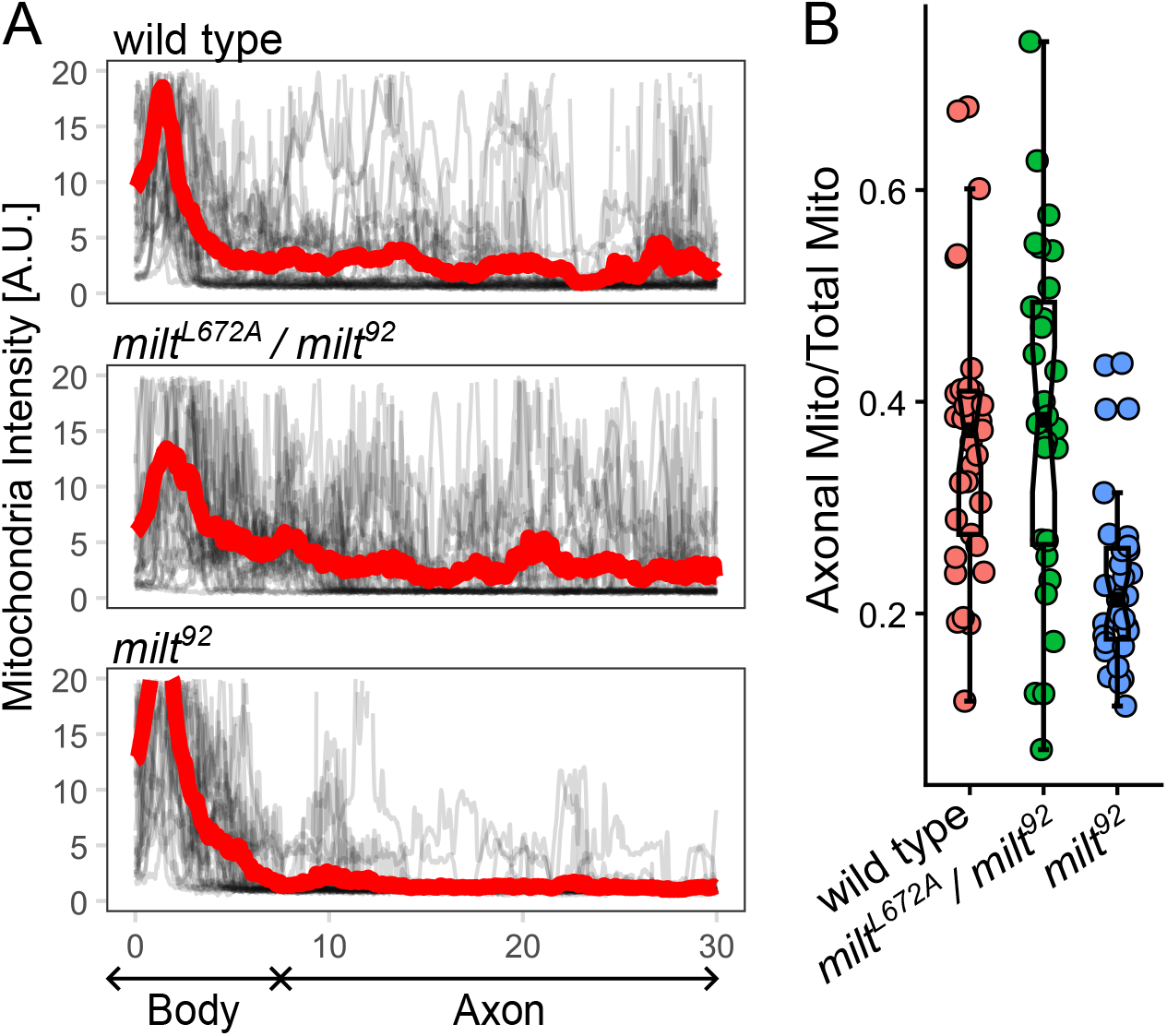
Mitochondrial distribution in hemizygous *milt*^*L672A*^ axons. A) Overlay of mitochondrial intensity distributions along axons of wild type, *milt*^*L672A*^ /*milt*^*92*^hemizygous and *milt*^*92*^ homozygous neurons. Grey lines represent individual cells and the red line indicates the average distribution for each genotype. B) Quantification of the axonal mitochondrial content defined as the fraction of total mitochondrial intensity located between 7.5 µm and 40 µm from the soma. Number of cells analysed; WT, 33; *milt*^*L672A*^ /*milt*^*92*^, 28; *milt*^*92*^, 31. Statistical significance was assessed using the Wilcoxon rank-sum test: *milt*^*92*^*p* differs from both WT and *milt*^*L672A*^ *(p* < 0.0002) whereas no significant difference is found between WT and *milt*^*L672A*^.

## Supplementary Videos

Supplementary videos 1 and 2: Negative gravitaxis of 3 day old and 38 day old flies, respectively. Genotype of flies is *Milt/milt*^*L672A*^ (left) and *milt*^*L672A*^*/milt*^*L672A*^ (right).

